# Regulation of (p)ppGpp hydrolysis by a conserved archetypal regulatory domain

**DOI:** 10.1101/257592

**Authors:** Séverin Ronneau, Julien Caballero-Montes, Aurélie Mayard, Abel Garcia-Pino, Régis Hallez

## Abstract

Sensory and regulatory domains allow bacteria to adequately respond to environmental changes. The regulatory ACT domains are mainly found in metabolic-related proteins as well as in long (p)ppGpp synthetase/hydrolase (SD/HD) enzymes. Here, we investigate the functional role of the ACT domain of SpoT, the only (p)ppGpp SD/HD of *Caulobacter crescentus*. We show that SpoT requires the ACT domain to hydrolyse ppGpp in an efficient way. In addition, our *in vivo* and *in vitro* data show that the phosphorylated version of EIIA^Ntr^ (EIIA^Ntr^~P) interacts directly with the ACT to inhibit the hydrolase activity of SpoT. Finally, we highlight the conservation of the ACT-dependent interaction between EIIA^Ntr^~P and SpoT/Rel along with the PTS^Ntr^-dependent regulation of (p)ppGpp accumulation upon nitrogen starvation in *Sinorhizobium meliloti*, a plant-associated α-proteobacterium. Thus, this work suggests that α-proteobacteria might have inherited from a common ancestor, a PTS^Ntr^ dedicated to modulate (p)ppGpp levels.

## Introduction

Bacteria use a wide range of sensory and regulatory domains to integrate environmental signals. In response to nutrient limitation, most bacteria synthesize a second messenger, the guanosine (penta) tetra-phosphate commonly referred to as (p)ppGpp. This molecule helps in reallocating cellular resources notably by reprograming transcription and interfering with cell cycle progression (Hallez et al., 2017; Hauryliuk et al., 2015). In *Escherichia coli*, (p)pppp levels are regulated by RelA and SpoT, two long RelA/SpoT homolog (RSH) enzymes. SpoT carries a synthetase domain (SD) that is poorly activated by starvation signals (carbon, fatty acids, phosphate, …) and a functional hydrolase domain (HD). By contrast, RelA harbours a SD activated by amino acids scarcity and a degenerated and inactive HD domain (reviewed in (Potrykus and Cashel, 2008) and (Hauryliuk et al., 2015)). Despite these differences, both RSH enzymes present the same domain architecture with catalytic domains (SD and HD) located towards the N-terminus (NTD) and regulatory domains (TGS for ThrRS, GTPase and SpoT and ACT for Aspartokinase, Chorismate mutase and TyrA) at the C-terminal end (CTD) (**Figure 1**). The CTD is thought to play a critical role in RSH by sensing the starvation signal and transducing it to the catalytic domains. RelA for example is known to be associated with the ribosome where it detects deacylated tRNA in the ribosomal acceptor site (A-site) as a signal for amino acid starvation, which activates SD activity (Haseltine and Block, 1973). Recent studies using cryo-electron microscopy revealed that *E. coli* RelA adopts an extended “open” conformation on stalled ribosomes with the A-site/tRNA in contact with the TGS domain in CTD (Arenz et al., 2016; Brown et al., 2016; Loveland et al., 2016). Likewise, activation of SpoT SD activity in *E. coli* cells starved for fatty acids has been shown to require an acyl carrier protein (ACP) that binds to the TGS of SpoT (Battesti and Bouveret, 2006). Although the role of CTD is undoubtedly critical for regulating catalytic functions of RSH, the mechanisms by which this regulation is mediated remain unknown.

**Figure 1.**
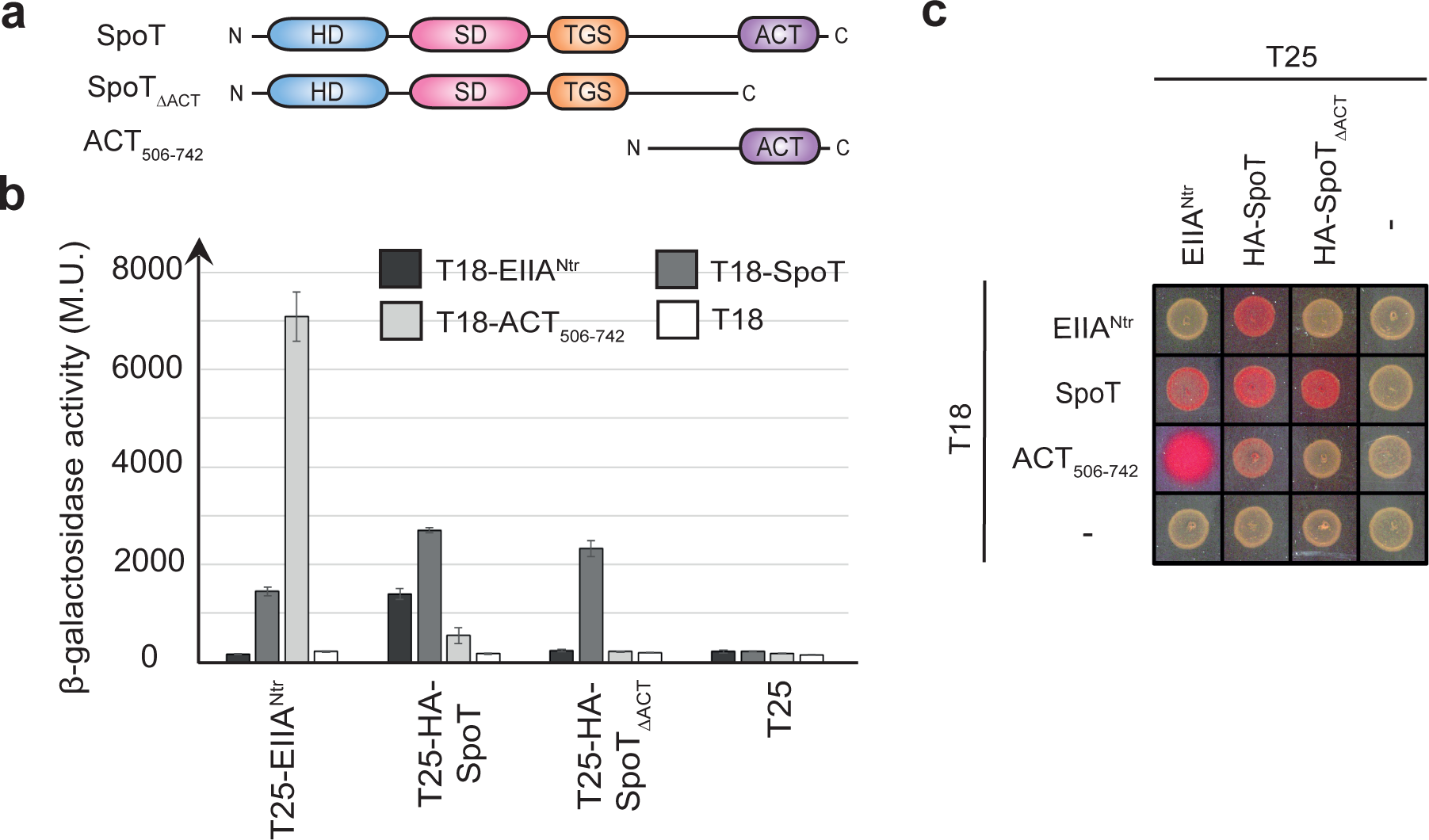
The ACT domain is required for the interaction between SpoT and EIIA^Ntr^~P in a bacterial two-hybrid assay. (a) Domain organization of *C. crescentus* SpoT with the catalytic domains, “hydrolase domain” (HD) and “synthetase domain” (SD), located at the N-terminal extremity and the regulatory domains, “ThrRS, GTPase and SpoT” (TGS) and “Aspartokinase, Chorismate mutase and TyrA” (ACT), located at the C-terminal end. (b) SpoT and ACT506-742, but not SpoT_ΔACT_, directly interact with EIIA^Ntr^~P. β-galactosidase assays were performed on MG1655 *cyaA::frt* (RH785) strains coexpressing T18-fused to *ptsN, spoT, ACT506-742* or alone with T25-fused to *ptsN, HA-spoT, HA-spoT*_ΔACT_ or alone. Error bars = SD, n = 3. The same strains were spotted on MacConkey Agar Base plates supplemented with 1% maltose. Plates were incubated for one day at 30 °C. The red color indicates positive interactions.

In contrast to *E. coli*, most α-proteobacteria encode bifunctional (SD/HD) RSH usually referred to as SpoT or Rel (Atkinson et al., 2011). This is the case of *Caulobacter crescentus* that harbours a single RSH (SpoT) sensitive to nitrogen or carbon starvation (Boutte and Crosson, 2011; Lesley and Shapiro, 2008). *C. crescentus* divides asymmetrically to generate two dissimilar progeny, a motile swarmer cell and a sessile stalked cell. The stalked cell initiates a new replication cycle (S phase) immediately at birth whereas the swarmer cell first enters into a non-replicative G1 phase before starting replication and concomitantly differentiating into a stalked cell (Curtis and Brun, 2010). Once accumulated, (p)ppGpp modulates cell cycle progression by specifically extending the G1/swarmer phase (Chiaverotti et al., 1981; Gonzalez and Collier, 2014). Recently, we reported a key role played by the nitrogen-related phosphotransferase system (PTS^Ntr^) in regulating (p)ppGpp accumulation in response to nitrogen starvation (Ronneau et al., 2016). We showed that EI^Ntr^, the first protein of PTS^Ntr^, uses intracellular glutamine concentration as a proxy for nitrogen availability, since glutamine binds to the GAF domain of EI^Ntr^ to inhibit its autophosphorylation. Therefore, glutamine deprivation strongly stimulates autophosphorylation of EI^Ntr^, which in turn triggers phosphorylation of the downstream components HPr and EIIA^Ntr^. Once phosphorylated, both HPr~P and EIIA^Ntr^~P modulate activities of SpoT to quickly increase (p)ppGpp levels. Whereas HPr~P stimulates SD activity of SpoT by an unknown mechanism, EIIA^Ntr^~P interacts directly with SpoT to interfere with its HD activity (Ronneau et al., 2016). The plant-associated α-proteobacterium *Sinorhizobium meliloti* also accumulates (p)ppGpp upon nitrogen or carbon starvation from a single RSH called Rel (Krol and Becker, 2011; Wells and Long, 2002), but the mechanism beyond this regulation remains unknown.

In this work, we investigate the role of the ACT domain in regulating the activity of RSH enzymes. In particular, we show that the ACT domain is indispensable for HD activity of *C. crescentus* SpoT and that EIIA^Ntr^~P inhibits (p)ppGpp hydrolysis by directly interacting with ACT of SpoT. In addition, we show that the EIIA^Ntr^~P-mediated regulation of RSH is conserved in S. *meliloti*, suggesting PTS^Ntr^ plays a critical role in sensing metabolic state and regulating cell cycle progression in α-proteobacteria.

## Results

### EIIA^Ntr^~P interacts directly with the ACT domain of SpoT

We showed previously that only the phosphorylated version of *C. crescentus* EIIA^Ntr^ (EIIA^Ntr^~P) was able to interact with SpoT in a bacterial two-hybrid (BTH) assay (Ronneau et al., 2016). To map the domains of SpoT interacting with EIIA^Ntr^~P, we used truncated versions of SpoT in a BTH assay (**Figure 1a**). We found that deleting the ACT domain (SpoT_ΔACT_) precluded the interaction with EIIA^Ntr^~P and with the isolated ACT domain (ACT_506-742_), but not with full-length SpoT (**Figure 1b-c**). Moreover, the ACT domain alone was able to strongly interact with EIIA^Ntr^~P (**Figure 1b-c**) as well as with itself (**Figure 1 – figure supplement 1**). Together, these results suggest that ACT-ACT interactions occur between SpoT subunits and show that the ACT domain is required to mediate the interaction between SpoT and EIIA^Ntr^~P.

### Inactivating the ACT domain increases (p)ppGpp levels

Disrupting the interaction between SpoT and EIIA^Ntr^~P should release the hydrolase (HD) activity of SpoT, since strains that either did not express EIIA^Ntr^ *(ΔptsN)* or expressed only the non-phosphorylated form of EIIA^Ntr^ (Δ*ptsP*, Δ*ptsH* or *ptsN*_*H66A*_) actively hydrolysed (p)ppGpp *in vivo* (Ronneau et al., 2016). Accordingly, a *C. crescentus* strain expressing *spoT*_*ΔACT*_ as the only copy of *spoT* should phenocopy the *ΔptsP* strain by decreasing motility and shortening G1 lifetime (Ronneau et al., 2016). Surprisingly, we found that *spoT*_*ΔACT*_ cells phenocopied rather the HD-dead mutant *spoT*_*D81G*_ than Δ*ptsP* (Ronneau et al., 2016) Indeed, the *spoT*_*ΔACT*_ mutation led to an increase of motility, an accumulation of G1/swarmer cells and a growth delay in PYE complex medium (**Figure 2a-d**). As for *spoT*_*D81G*_, these phenotypes might be due to a (p)ppGpp excess in *spoT*_*ΔACT*_. In support of this, *spoT*_*ΔACT*_ cells accumulated (p)ppGpp without stress (**Figure 2e**). In addition, abolishing the synthetase (SD) activity in *spoT*_*ΔACT*_ cells, either by deleting *ptsP* (EI^Ntr^) or by incorporating a Y323A substitution into SpoT (*spoT*_*Y323A ΔACT*_) known to specifically inactivate SD activity (Boutte and Crosson, 2011; Ronneau et al., 2016), suppressed all the phenotypes (**Figure 2a-e and Figure 2 – figure supplement 1a**). The (p)ppGpp accumulation observed in *spoT*_*ΔACT*_ cells could be due to an increase of SD activity or a decrease of HD activity. To discriminate between these two possibilities, we measured (p)ppGpp levels in nitrogen-deplete (-N) conditions, which mainly relies on SD (Ronneau et al., 2016). We observed that *spoT*_*ΔACT*_ and SpoT_D81G_ protein levels in the corresponding mutant strains were slightly higher than the SpoT level in the wild-type strain (**Figure 2 – figure supplement 1b-c**). Nevertheless, upon nitrogen starvation, neither *spoT*_*ΔACT*_ nor *spoT*_*D8IG*_ cells produced more (p)ppGpp than wild-type cells (**Figure 2 – figure supplement 1d**). This suggests that SD activity is not enhanced in the of *spoT*_*ΔACT*_ nor *spoT*_*D8IG*_ strains. The slight increase of *spoT*_*ΔACT*_ nor *spoT*_*D8IG*_ levels is likely a result of the positive feedback loop of (p)ppGpp on *spoT* promoter. Indeed, *P*_*spoT*_ displayed higher activity in strains accumulating (p)ppGpp (*ptsP*_*L83Q*_ and *P*_*xylx*_::*relA-FLAG*) and lower activity in ppGpp^0^ strains (*ΔptsP, ΔptsH and ΔptsN*), so that SpoT protein levels varied according to (p)ppGpp levels (**Figure 2 – figure supplement 1e-g**). Thus, the higher concentration of (p)ppGpp detected in *spor*_*ΔACT*_ cells may come from an inactivation of HD activity, thereby suggesting that ACT is required for HD activity.

**Figure 2.**
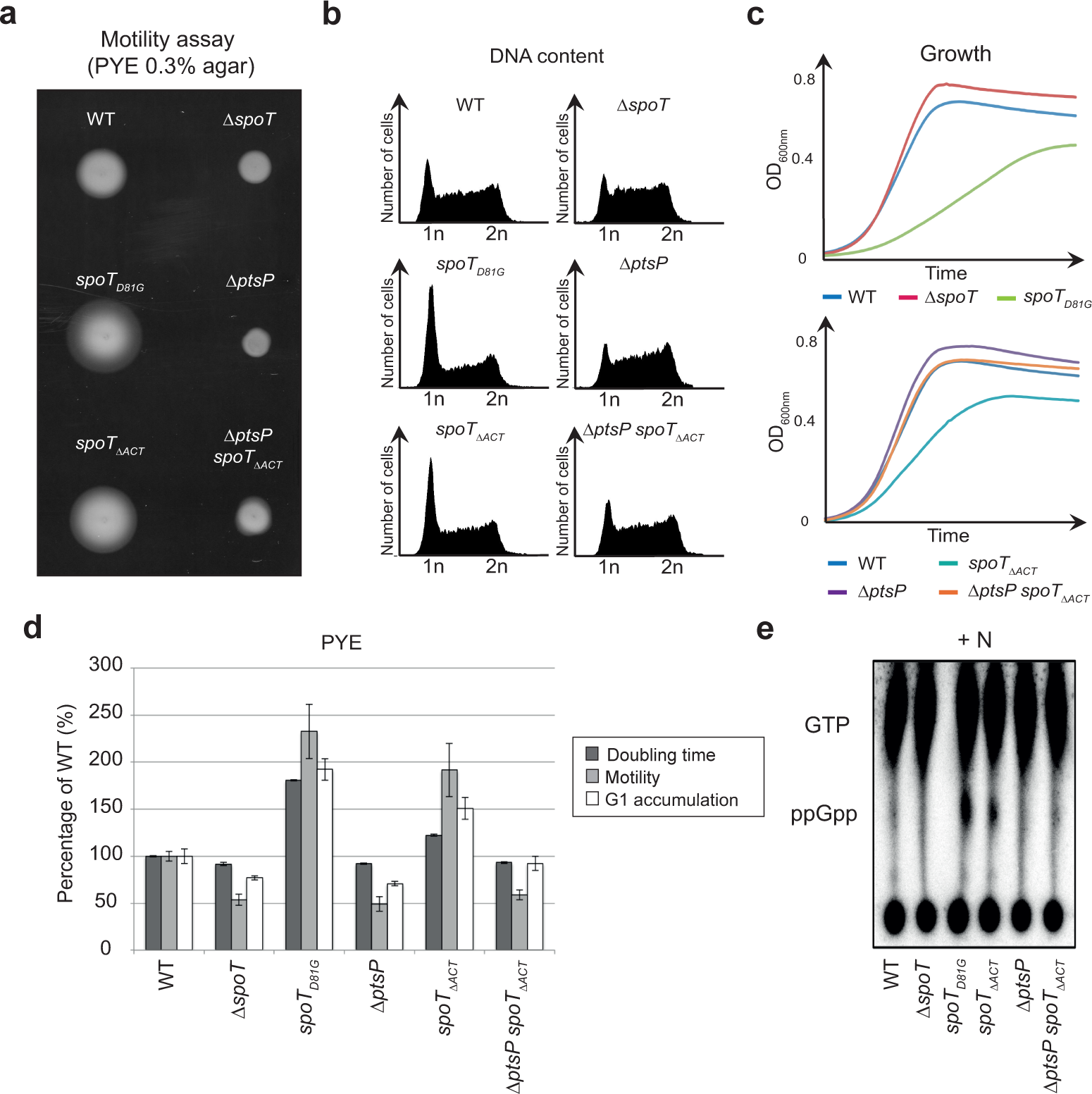
Deletion of the ACT domain of SpoT leads to (p)ppGpp accumulation, which consequently extends the G1/swarmer cell lifetime of *C. crescentus*. (a-d) Extension of the G1/swarmer lifetime in *spoT*_*ΔACT*_ cells is suppressed by the deletion of *ptsP* (encoding EI^Ntr^). Motility (a), DNA content (b) and growth (c) were measured in WT (RH50), *ΔspoT* (RH1755), *spoT*_*D81G*_ (RH1752), *ΔptsP* (RH1758), *spoT*_ΔACT_ (RH1476) and *ΔPtsp spoT*_ΔACT_ (RH1478) grown in complex media (PYE). (d) Data shown in a-c were normalized to the WT (100%). Error bars = SD, n = 3. (e) Deletion of the ACT domain leads to (p)ppGpp accumulation. The intracellular levels of (p)ppGpp were evaluated by TLC after nucleotides extraction from the same strains grown in nitrogen-replete (+N) conditions.

### The hydrolase activity of SpoT requires the ACT domain

To test whether ACT was required for HD activity, we first measured the endogenous HD activity of SpoT *in vivo* (Ronneau et al., 2016). To this end, we used *C. crescentus* strains in which (p)ppGpp (i) cannot be produced anymore by the endogenous SpoT since its SD activity has been inactivated with the Y323A mutation (Boutte and Crosson, 2011), but (ii) can be synthesized upon addition of xylose by a truncated version of the *E. coli* RelA (p)ppGpp synthetase expressed from the xylose-inducible promoter (*P*_*xyIx*_::*relA-FLAG*) at the *xylX* locus (Gonzalez and Collier, 2014). Thus in these strains the only HD activity capable to degrade (p)ppGpp produced by *E. coli* RelA was supplied by endogenous SpoT variants. In agreement with our previous observations, inactivation of SpoT HD in such a background (*spoT*_*D81G y323A*_: P _*xyIx*_::*relA-FLAG*,) led to a strong (p)ppGpp accumulation, an extension of the G1 phase and a growth arrest (**Figure 3**; (Ronneau et al., 2016)). In contrast, releasing SpoT HD activity (*ΔptsP spoT*_*Y323A*_: P _*xyIx*_::*relA-FLAG*) led to undetectable levels of (p)ppGpp, a reduced G1 phase and an optimal growth (Figure 3; (Ronneau et al., 2016)). Interestingly, deleting the ACT domain of SpoT (*spoT*_*Y323A ΔACT*_,· P_*xylx*_::*relA-FLAG*) led to a (p)ppGpp accumulation, a G1 proportion and a growth rate similar to the HD-dead mutant (*spoT*_*D81G y323A*_: P_*xylx*_::*relA-FLAG*), and this independently of the presence of *ptsP* (**Figure 3**).

**Figure 3.**
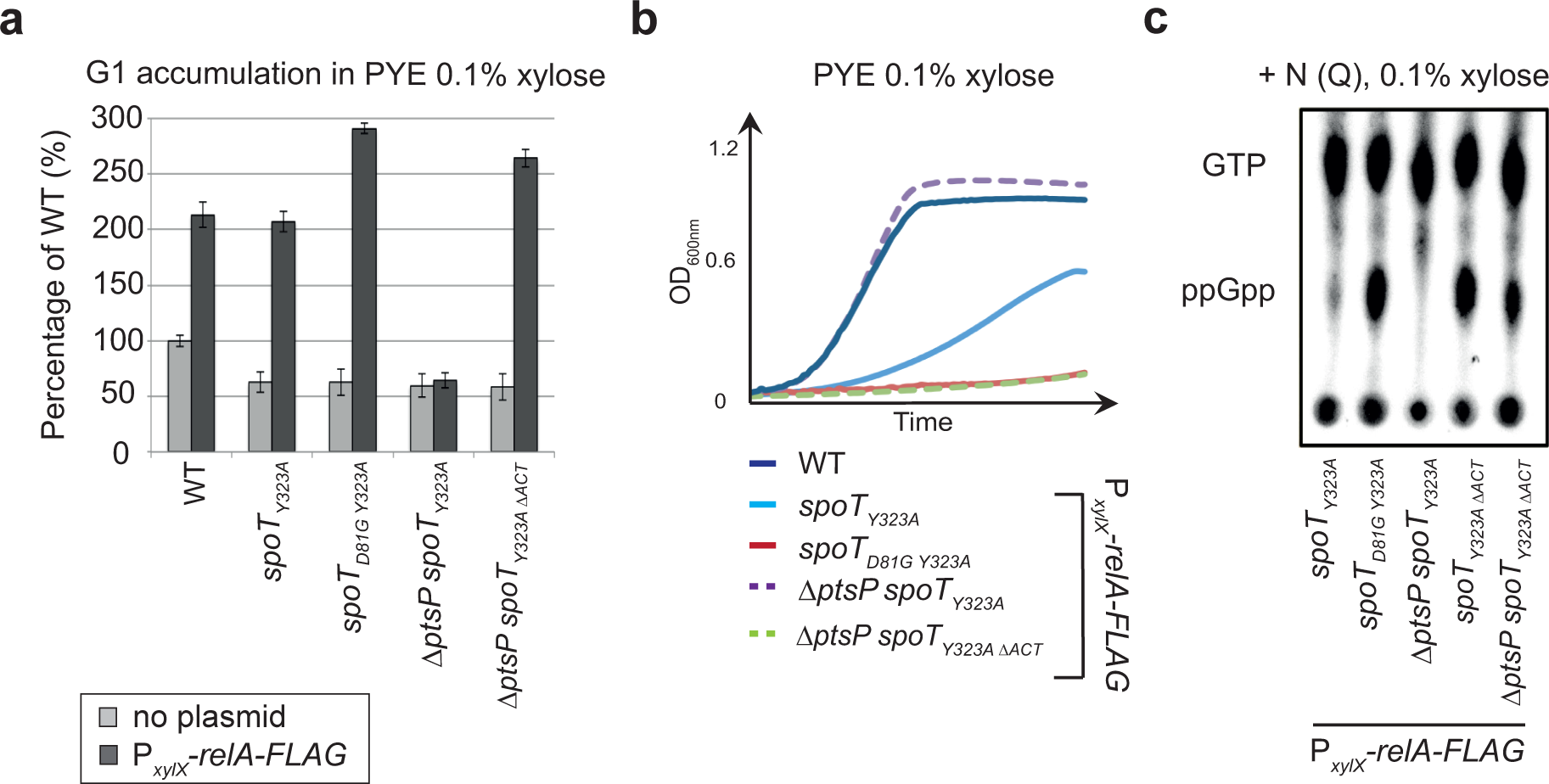
The ACT domain is required *in vivo* to support the hydrolase activity of SpoT. (a-b) Absence of the ACT domain of SpoT promotes G1 accumulation and decreases growth rate upon artificial exogenous production of (p)ppGpp. (a) Flow cytometry analysis to determine DNA content in asynchronous population and (b) growth of WT (RH50), *spoT*_*Y323A*_ (RH1844), *spoT*_*D81G Y323A*_ (RH2193), *ΔptsP spoTY323A* (RH2196) and *ΔptsP spoT* _*Y323A ΔACT*_ (RH2491) with (black bars) or without (grey bars) *P*_*xylx*_::*relA-FLAG* in PYE medium supplemented with 0.1% of xylose. The flow cytometry data were normalized to the WT without *P*_*xylX*_::*relA-FLAG* (100%). Error bars = SD, n = 3. (c) The regulatory ACT domain of SpoT is required *in vivo* to degrade (p)ppGpp in nitrogen-replete condition (+N). The intracellular levels of (p)ppGpp were evaluated by TLC after nucleotides extraction from *spor*_*Y323A*_ (RH1844), *spoT_D81G Y323A_* (RH2193), *ΔptsP spoT*_*Y323A*_ (RH1586), *spoT*_*Y323A ΔACT*_ (RH2491) and *ΔptsP spoT*_*Y323A ΔACT*_ (RH2491) harbouring *P*_*xylx*_::*relA-FLAG* and grown in nitrogen-replete (+N) media supplemented with 0.1% xylose.

These data strongly suggest that the ACT domain is strictly required *in vivo* for the hydrolase activity of SpoT and that EIIA^Ntr^~P inhibits HD activity of SpoT by interfering with the ACT domain. To test these hypotheses, we performed *in vitro* HD assays with the full-length enzyme and mutants lacking the ACT domain (SpoT_ΔACT_) or containing only the two catalytic domains (SpoT_1-373_). We observed that full-length SpoT could efficiently degrade ^[32P]^ppGpp but the deletion of the ACT domain in both SpoT_ΔACT_ and SpoT1-373 strongly affected HD activity, since radiolabelled ^[32P]^ppGpp remained mostly intact in the presence of both ACT-deficient SpoT variants (**Figure 4a**, compare lane 3 to lanes 4-7). To check whether phosphorylated EIIA^Ntr^ could modulate the HD activity of SpoT, ^[32P]^ppGpp hydrolysis was measured in the presence of EIIA^Ntr^ and non-phosphorylatable (EIIA^Ntr^_H66A_) or phosphomimetic (EIIA^Ntr^_H66E_) version of EIIA^Ntr^. We found that only EIIA^Ntr^_H66E_ was able to interfere with SpoT HD activity (**Figure 4b-c**). The role of phosphorylation was further confirmed using EIIA^Ntr^~P. Our results show that EIIA^Ntr^~P efficiently inhibited SpoT HD activity, protecting ^[32P]^ppGpp from hydrolysis (**Figure 4d and Figure 4 – figure supplement 1**). Altogether, these data show that EIIA^Ntr^~P inhibits the HD activity of SpoT by directly interacting and likely interfering with the ACT domain.

**Figure 4.**
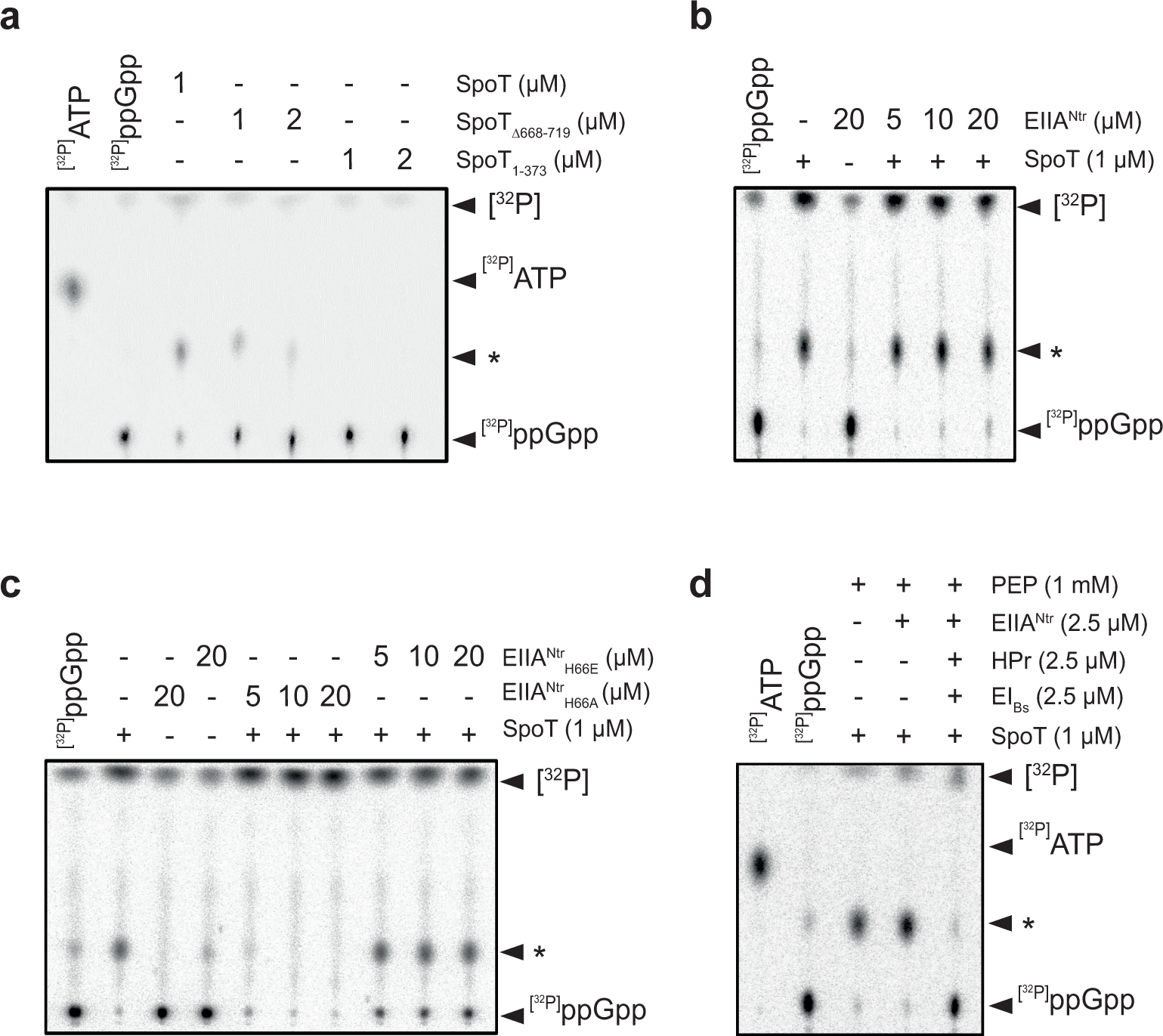
EIIA^Ntr^~P specifically inhibits the hydrolase activity of SpoT *in vitro* in an ACT-dependent way. (a) The ACT domain of SpoT is required to hydrolyse ppGpp. Radiolabelled ^[32P]^ppGpp incubated with 1 *µ*M SpoT, 1 *μ*M or 2 *μ*MSpoT_Δ668-719_ or SpoT1-373 was separated by TLC. (b-c) Phosphomimetic variant of EIIA^Ntr^ inhibits the hydrolase activity of SpoT. Radiolabelled ^[32P]^ppGpp incubated with 1 *μ*M SpoT in the presence of increasing concentrations (5, 10 and 20 *μ*M) of unphosphorylated EIIA^Ntr^ (b), non-phosphorylatable or phosphomimetic variants of EIIA^Ntr^ (c) was separated by TLC. ^[32P]^ppGpp incubated with EIIA^Ntr^ variants alone were added as controls. (d) Phosphorylated EIIA^Ntr^ fully protects ppGpp from hydrolysis by SpoT. Radiolabelled ^[32P]^ppGpp incubated with 1 SpoT in the presence of 2.5 EIIA^Ntr^ with of without EIbs (2.5 *μ*M) and HPr (2.5 *μ*M) was separated by TLC. The star “*” indicates a ^[32P]^ppGpp intermediate degradation. ^[32P]^ATP and ^[32P]^ppGpp were used as references.

### PTS^Ntr^ regulates (p)ppGpp accumulation in *Sinorhizobium meliloti*

The inhibition of EI^Ntr^ autophosphorylation by glutamine was first observed in the γ-proteobacterium *Escherichia coli* (Lee et al., 2013) and the plant-associated α-proteobacterium *Sinorhizobium meliloti* (Goodwin and Gage, 2014). In addition, S. *meliloti* also accumulates (p)ppGpp in response to nitrogen starvation (Krol and Becker, 2011; Wells and Long, 2002). This suggests that PTS^Ntr^ stimulates (p)ppGpp accumulation in response to glutamine deprivation as shown for *C. crescentus* (Ronneau et al., 2016). To test this hypothesis, we checked whether the nitrogen-related EIIA component of S. *meliloti* (EIIA^Ntr^) was able to interact with the bifunctional (SD/HD) RSH of S. *meliloti* (Rel) in a BTH assay. As shown in **Figure 5a**, the full-length Rel fused to T25 (T25-Rel) interacted with EIIA^Ntr^ fused to T18 (T18-EIIA^Ntr^). Similarly to *C. crescentus*, the deletion of ACT (T25-Rel_ΔACT_) abolished this interaction (Figure 5a). Another conserved feature is that phosphorylation of EIIA^Ntr^ enhanced interaction with Rel. Indeed, we previously showed that (i) EIIA^Ntr^ was phosphorylated in the BTH assay by the endogenous PTS^Ntr^ system of *E. coli* (EI^Ntr^ and NPr) and (ii) the interaction between SpoT and EIIA^Ntr^ was altered in an *E. coli Anpr* background (Ronneau et al., 2016). Likewise, the interaction between S. *meliloti* T18-EIIA^Ntr^ and T25-Rel was strongly diminished in a Δnpr background (**Figure 5b**). This indicates that phosphorylation of EIIA^Ntr^ also enhances the interaction with Rel. We constructed single in-frame deletion of *ptsP* (SMc02437) and *rel* (SMc02659) genes to test the PTS^Ntr^-dependent accumulation of (p)ppGpp in S. *meliloti* in response to nitrogen deprivation. Upon nitrogen starvation (-N), neither *Δrel* nor *ΔptsP* cells accumulated (p)ppGpp in contrast to wild-type cells (**Figure 5c**). The increase of G1 proportion that results from the accumulation of (p)ppGpp upon nutrient limitation was previously used to synchronize S. *meliloti* (De Nisco et al., 2014). Thus if PTS^Ntr^ regulates (p)ppGpp levels, we reasoned that the G1 proportion of Δ*rel* and Δ*ptsP* populations should be reduced. This is exactly what we observed, with a proportion of G1 cells in the *ΔptsP* and Δ*rel* strains reduced in comparison to the wild-type strain (**Figure 5d-e**). In contrast, the ectopic production of (p)ppGpp in non-starved S. *meliloti* cells with an IPTG-inducible version of RelA_Ec_ strongly increased the proportion of G1 cells and led to growth arrest (**Figure 5 – figure supplement 1**). Altogether, these results support a conserved role of PTS^Ntr^ in regulating (p)ppGpp accumulation in response to nitrogen starvation as well as in the control of cell cycle progression in S. *meliloti*.

**Figure 5.**
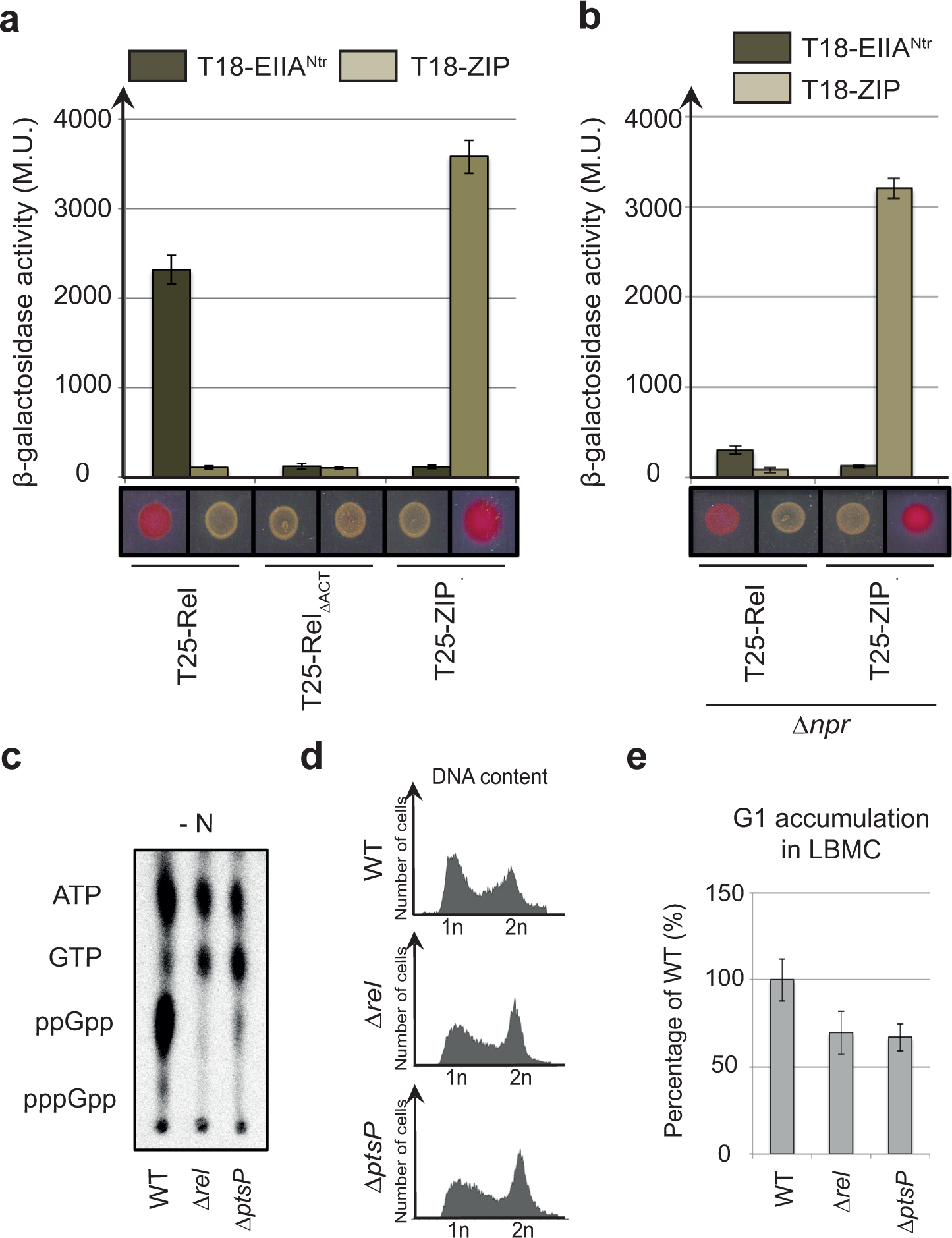
PTS^Ntr^ modulates (p)ppGpp accumulation and cell cycle progression in *Sinorhizobium meliloti*. (a-b) EIIA^Ntr^ interacts with Rel in a ACT-dependent way. β-galactosidase assays were performed on (a) MG1655 *cyaA::frt* (RH785) or (b) MG1655 *cyaA::frt Anpr* (RH2122) strains coexpressing T18-fused to *ptsN* or *ZIP* with T25-fused to *rel, rel*_*ΔACT*_ or *ZIP*. Error bars = SD, n = 3. The same strains were spotted on MacConkey Agar Base plates supplemented with 1% maltose. Plates were incubated for one day at 30 °C. The red color indicates positive interactions. (c-e) A functional PTS^Ntr^ is required for (p)ppGpp accumulation in S. *meliloti* upon nitrogen starvation (-N) and for cell cycle progression in nitrogen-replete (+N) condition. (c) The intracellular levels of (p)ppGpp were evaluated by TLC after nucleotides extraction from WT (RH2000), *ΔrelA* (RH2327) and *ΔptsP* (RH2326) grown in nitrogen-deplete (-N) conditions. (d-e) DNA content (d) and G1 proportion were measured in the same strains grown in complex media (LBMC). G1 proportions were normalized to the WT (100%). Error bars = SD, n = 3.

## Discussion

Two decades ago, mutational analysis of the *spoT* gene of *E. coli* suggested that the C-terminal end of SpoT including the ACT domain was required to bind regulators of the hydrolase activity of SpoT (Gentry and Cashel, 1996). Later, Mechold et al. reported that deleting the C-terminal domain of SpoT in *Streptococcus equisimilis* led to a strong inhibition of (p)ppGpp degradation *in vitro* of about 150 fold in comparison to the full-length protein (Mechold et al., 2002). In agreement with these studies, our work supports that the regulatory ACT domain modulates SpoT hydrolase (HD) activity in *C. crescentus*. In a nitrogen-rich environment (+N), the ACT domain is free to stimulate HD activity, thereby limiting (p)ppGpp concentration. Upon nitrogen starvation (-N), the last component of the nitrogen-related PTS (PTS^Ntr^), EIIA^Ntr^, is phosphorylated and binds the ACT domain of SpoT. This physical interaction interferes with the ACT-dependent stimulation of SpoT HD activity (**Figure 6**). The recent RelA structures on stalled ribosomes highlighted the role of C-terminal domain (CTD) in sensing nutrient availability (Arenz et al., 2016; Brown et al., 2016; Loveland et al., 2016). When bound to the ribosome, RelA adopts an “open” conformation, which is thought to relieve the inhibitory effect of CTD on its synthetase (SD) activity and leads to (p)ppGpp synthesis (Gropp et al., 2001). This “open” conformation seems to be favoured by specific interactions between the ribosome stalk, the A-site finger and the tRNAs with the CTD (Loveland et al., 2016). Given the conserved domain architecture of RelA and SpoT proteins, we propose that EIIA^Ntr^~P modulates SpoT conformation to decrease its HD activity (**Figure 6**). Since ACT seems to interact with itself, it is tempting to speculate that dimerization of ACT induces a conformation that enhances the HD activity. Oligomerization of long RSH has already been proposed to regulate their activity (Gropp et al., 2001; Jain et al., 2006). In such a scenario, EIIA^Ntr^~P would preclude or interfere with the active conformational state. The ACT domain is highly prevalent among RSH enzymes, thus the role of the ACT domain in sustaining the hydrolase activity in these RSH is likely conserved as well.

**Figure 6.**
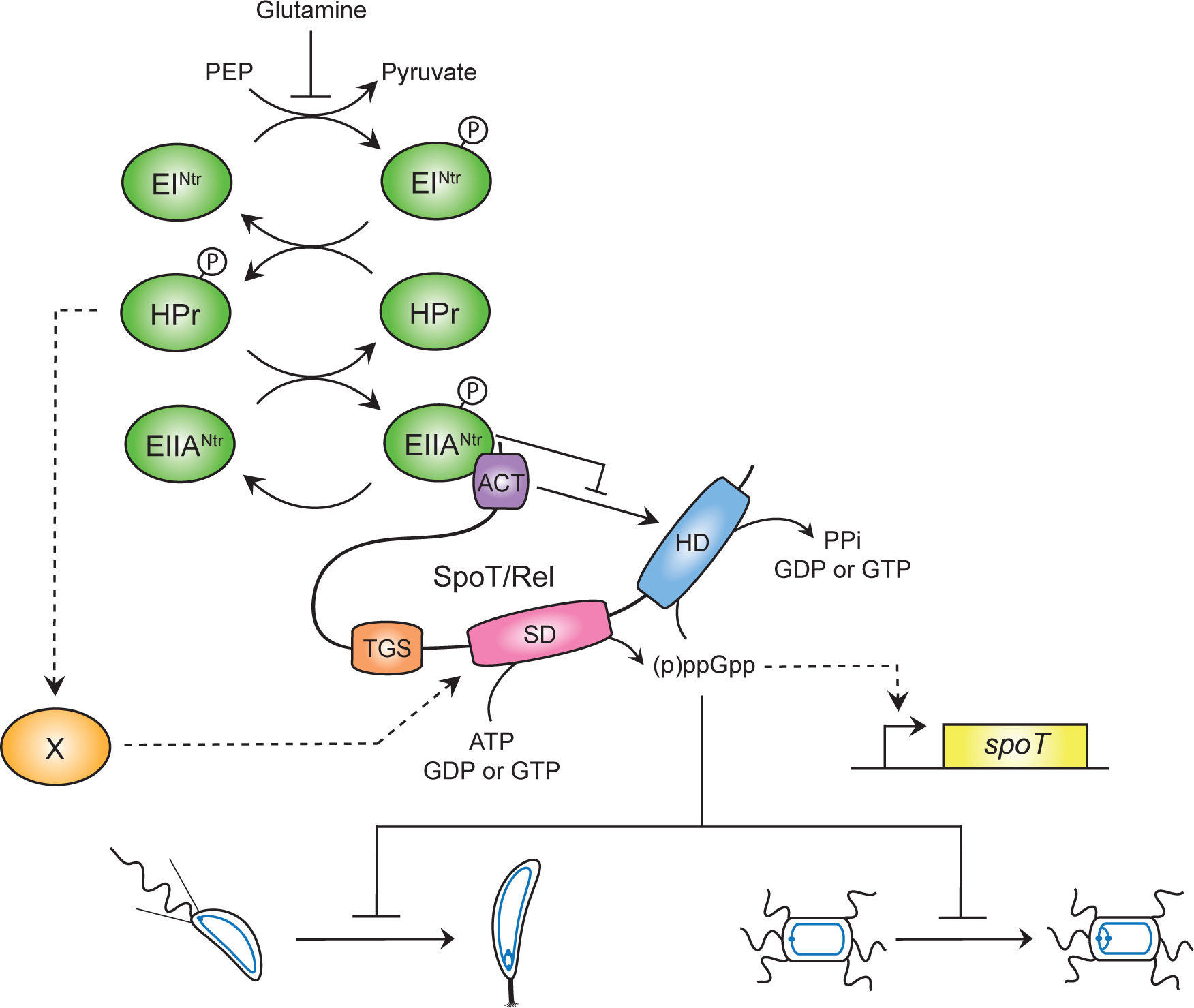
EIIA^Ntr^~P binds the ACT domain of SpoT to inhibit its hydrolase activity (HD). Upon nitrogen starvation (*i.e*. glutamine deprivation), EI^Ntr^ phosphorylates its downstream components, HPr and EIIA^Ntr^. As a consequence, EIIA^Ntr^~P interacts with ACT to inhibit its stimulating effect on the hydrolase activity (HD) of SpoT. This inhibition avoids to degrade (p)ppGpp produced by the synthetase domain (SD), this latter being strongly stimulated by HPr~P by an unknown mechanism. In addition, increasing the (p)ppGpp level promotes the accumulation of the SpoT protein by a positive feedback loop mechanism on transcription of the *spoT* gene. Ultimately, the burst of (p)ppGpp delays the G1-to-S transition of *C. crescentus* (left) and S. *meliloti* (right) cells.

We also reported that SpoT_ΔACT_ is still able to increase (p)ppGpp concentration upon nitrogen starvation by stimulating SD activity in a PTS^Ntr^-dependent way (**Figure 2 – figure supplement 1d**). This regulation could involve the TGS domain as described for *E. coli*, in which stimulation of SpoT SD activity upon fatty acid starvation depends on the TGS domain (Battesti and Bouveret, 2006). In support of this, deletion of the CTD harboring the TGS and the ACT domains completely abolished the ability of *C. crescentus* SpoT to produce (p)ppGpp upon nitrogen starvation (Boutte and Crosson, 2011). Altogether, our data support that regulators bind to SpoT via the regulatory CTD domains to modulate enzymatic activities.

Besides *C. crescentus*, another α-proteobacterium, *Sinorhizobium meliloti*, uses PTS^Ntr^ to sense nitrogen starvation and to stimulate (p)ppGpp accumulation (**Figure 6**). We found that EIIA^Ntr^ interacts with Rel (**Figure 5a**) and that phosphorylation of EIIA^Ntr^ promotes this interaction (**Figure 5b**). As glutamine also inhibits phosphorylation of EI^Ntr^ in S. *meliloti* (Goodwin and Gage, 2014), glutamine deprivation likely leads to hyperphosphorylation of downstream PTS^Ntr^ components, favouring subsequent interaction between EIIA^Ntr^~P and Rel. In support of that, we showed S. *meliloti* required EI^Ntr^ protein to accumulate (p)ppGpp upon nitrogen starvation (**Figure 5c**), as previously shown for *C. crescentus* (Ronneau et al., 2016). In addition, we observed that the physical interaction between Rel and the phosphorylated form of EIIA^Ntr^ also occurs in another α-proteobacterium, *Rhodobacter sphaeroides* (**Figure 5 – figure supplement 2**). Altogether, our data support that the PTS^Ntr^-dependent control of SpoT/Rel activities is another conserved feature in α-proteobacteria (Hallez et al., 2004).

Similarly to *C. crescentus* (Gonzalez and Collier, 2014; Ronneau et al., 2016), we found that (p)ppGpp accumulation also delays the G1-to-S transition in S. *meliloti*, (**Figure 5d-e and Figure 5 – figure supplement 2**). In fact, this feature has been used to synchronize a population of S. *meliloti* by transiently blocking bacteria starved for carbon and nitrogen in G1 phase of the cell cycle (De Nisco et al., 2014). The specific interference of (p)ppGpp with the G1-to-S transition of the cell cycle could be another conserved feature in α-proteobacteria.

## Methods

### Bacterial strains and growth conditions

Oligonucleotides, strains and plasmids used in this study are listed in **Supplementary Table. 1, 2 and 3**, together with construction details provided in the Supplementary Methods. *Escherichia coli* Top10 was used for cloning purpose, and grown aerobically in Luria-Bertani (LB) broth (Invitrogen) (Casadaban and Cohen, 1980). Electrocompetent cells were used for transformation of *E. coli*. All *Caulobacter crescentus* strains used in this study are derived from the synchronizable (NA1000) wild-type strain, and were grown in Peptone Yeast Extract (PYE) or synthetic M2 (20 mM PO_4_^3-^, 9.3 mM NH_4_^+^; +N) or P2 (20 mM PO_4_^3-^; -N) supplemented with 0.5 mM MgSO_4_, 0.5 mM CaCl_2_, 0.01 mM FeSO_4_ and 0.2% glucose (respectively M2G or P2G) media at 28-30 °C. Growth was monitored by following the optical density at 660 nm (OD660) during 24 hrs, in an automated plate reader (Epoch 2, Biotek) with continuous shaking at 30 °C. Motility was monitored on PYE swarm (0.3 % agar) plates. Area of the swarm colonies were quantified with ImageJ software as described previously (Ronneau et al., 2016). For kinetic experiments with *Sinorhizobium meliloti* (**Figure 5 – figure supplement 1d**), bacteria where cultivated overnight at 30 °C in LBMC (LB broth with 2.5 mM MgSO_4_ and 2.5 mM CaCl_2_) supplemented with kanamycin, then back-diluted in LBMC during 3 hrs before induction with 0.1 mM IPTG. Samples were taken each hour during 5 hrs. For *E. coli*, antibiotics were used at the following concentrations µg/ml; in liquid/solid medium): ampicillin (50/100), kanamycin (30/50), oxytetracycline (12.5/12.5) where appropriate. For *C. crescentus*, media were supplemented with kanamycin (5/20), tetracycline (1/2.5) where appropriate. The doubling time of *Caulobacter* strains was calculated in exponential phase (OD660: 0.2 - 0.5) using D = [ln(2).(T_(B)_ – T_(A)_)) / (ln(OD_660(B)_)-ln(OD_660(A)_)] and normalized according to the wild-type strain. *E. coli* S17-1 and *E. coli* MT607 helper strains were used for transferring plasmids to *C. crescentus* by respectively bi- and tri-parental mating. In-frame deletions were created by using pNPTS138-derivative plasmids and by following the procedure described previously (Ronneau et al., 2016).

### Bacterial two-hybrid assays

Bacterial two-hybrid (BTH) assays were performed as described previously in (Ronneau et al., 2016). Briefly, 2 μl of MG1655 *cyaA::frt* (RH785) and MG1655 *cyaA::frt Δnpr* (RH2122) strains expressing T18 and T25 fusions were spotted on MacConkey Agar Base plates supplemented with ampicillin, kanamycin, maltose (1%), and incubed for 1 day at 30 °C. All proteins were fused to T25 (pKT25) or T18 (pUT18C) at their N-terminal extremity. The β-galactosidase assays were performed as described in (Ronneau et al., 2016). Briefly, 50 μl *E. coli* BTH strains cultivated overnight at 30° C in LB medium supplemented with kanamycin, ampicillin and IPTG (1 mM) were resuspended in 800 μl of Z buffer (60 mM Na_2_HPO_4_, 40 mM NaH_2_PO_4_, 10 mM KCl, 1 mM MgSO_4_) and lysed with chloroform. After the addition of 200 μl ONPG (4 mg/ml), reactions were incubated at 30° C until color turned yellowish. Reactions were then stopped by adding 500 μl of 1 M Na2CO3, and absorbance at 420 nm was measured. Miller Units are defined as (OD420 × 1,000)/ (OD590 × t × v), where “ OD590” is the absorbance of the cultures at 590 nm before the β-galactosidase assays, “t” is the time of the reaction (min), and “v” is the volume of cultures used in the assays (ml). All the experiments were performed with at least three biological replicates.

### Flow cytometry analysis

DNA content was measured using Fluorescence-Activated Cell Sorting (FACS) as described previously in (Ronneau et al., 2016). Briefly, cells were fixed in ice-cold 70% Ethanol. Fixed samples were then washed twice in FACS staining buffer (10 mM Tris pH 7.2, 1 mM EDTA, 50 mM NaCitrate, 0.01% Triton X-100) containing 0.1 mg/ml RNaseA and incubated at room temperature (RT) for 30 min. Cells were then harvested by centrifugation for 2 min at 8,000 x g, resuspended in 1 ml FACS staining buffer containing 0.5 μM Sytox Green Nucleic acid stain (Life Technologies), and incubated at RT in the dark for 5 min. Samples were analyzed in flow cytometer (FACS Calibur, BD Biosciences) at laser excitation of 488 nm. Percentage of gated G1 cells of each strain was then normalized using gated G1 cells of the wild-type strain as reference.

### Detection of intracellular (p)ppGpp levels

(p)ppGpp levels were visualized as described previously in (Ronneau et al., 2016) for *Caulobacter crescentus* (Cc) and in (Krol and Becker, 2011; Wells and Long, 2002) for *Sinorhizobium meliloti* (Sm). Briefly, strains were grown overnight in PYE and then diluted for a second overnight culture in M5GG (Cc) or grown overnight in LBMC medium (Sm). Then, cells were diluted a second time in M5GG (Cc) or in LBMC medium (Sm) and grown for 3 hrs to reach an OD660 of 0.5 (Cc) or 0.7 (Sm). Cells were then split into two parts and washed twice with P5G-labelling buffer (Cc; (Ronneau et al., 2016)) or with MOPS-MGS without glutamate (Sm; (Mendrygal and Gonzalez, 2000)). One milliliter of cells were then resuspended in 225 μl of P5G-labelling (-N) or M5G-labelling (+N) for *C. crescentus* and in 225 μl of MOPS-MGS with (+N) or without (-N) glutamate and 0.05% NH_4_^+^ for S. *meliloti*. In addition, media were supplemented with 25 μl of KH_2_^32^PO_4_ at 100 μCi ml^-1^ and incubated for 1 hr (Sm) or 2 hrs (Cc) with shaking at 30 °C. Then, samples were extracted with an equal volume of 2 M formic acid, placed on ice for 20 min and then stored overnight at -20 °C. All cell extracts were pelleted at 14,000 rpm (18,000 x g) for 3 min and 6 x 2 μl (Cc) or 3 x 2 ill (Sm) of supernatant were spotted onto a polyethyleneimine (PEI) plate (Macherey-Nagel). PEI plates were then developed in 1.5 M KH_2_PO_4_ (pH 3.4) at room temperature. Finally, TLC plates were imaged on a MS Storage Phosphor Screen (GE Healthcare) and analysed with Cyclone Phosphor Imager (PerkinElmer). For hydrolase experiments (Figure 3C), cells were incubated 1 hr in P5G supplemented with xylose (0.1%). Then, cells were washed twice with P5G-labelling and resuspended in P5G-labelling (-N) supplemented with KH_2_^32^PO_4_, xylose (0.1%) and glutamine (9.3 mM).

### β-galactosidase assay

The β-galactosidase assays performed to measure *P*_*spoT*_-*lacZ* activity were essentiality done as for the BTH assay with the following modifications. One milliter of *Caulobacter* strains harbouring the *P*_*spoT*_-*lacZ* fusion was resuspended in 800 μl of Z buffer and the absorbance of the cultures at 660 nm (OD660) instead of 590 nm was measured before the β-galactosidase assays. All the experiments were performed with three biological replicates and were normalized according to the wild-type strain harbouring the *P*_*spoT*_-*lacZ* fusion cultivated at 30°C.

### Immunoblot analysis

Immunoblot analyses were performed as described in (Beaufay et al., 2015) with the following primary antibodies: anti-MreB (1:5,000) (Beaufay et al., 2015), anti-SpoT (1:5,000) and secondary antibodies: anti-rabbit linked to peroxidase (GE Healthcare) at 1:5,000, and visualized thanks to Western Lightning Plus-ECL chemiluminescence reagent (Biorad) and ImageQuant LAS400 (GE Healthcare).

### Proteins purification

The purification of *B. subtilis* EI was essentially done as described in (Galinier et al., 1997), with the modifications indicated below. Genes encoding *C. crescentus* SpoT, SpoT_D81G_, SpoT _1-373_, SpoT_ΔAcτ_, as well as HPr, EIIA^Ntr^, EIIA^Ntr^_H66E_ and EIIA^Ntr^_H66A_ were transformed into *E. coli* BL21 (DE3) for protein production. In the case of *B. subtilis* EI, *E. coli* NM522 was transformed with plasmid pQE-30-*ptsI_Bs* as described in (Galinier et al., 1997) for protein expression.

Each protein contains an N-terminal 6His-tag for purification by Ni-NTA chromatography. Cells were grown to an OD_600_ of ~ 0.7 in LB medium (1 L) at 37 °C. Isopropyl-β-D-thiogalactoside (IPTG) was added to a final concentration of 0.5 mM and incubated overnight at 28 °C. Cells were harvested by centrifugation for 20 min at 6,000 x g, 4°C, resuspended in KCl 8 mM, TCEP 1 mM, MgCl_2_ 2 mM, Tris 50 mM at pH 8, Protease Inhibitor Cocktail (Roche) and lysed with a cell disruptor at 60 psi in KCl 500 mM, TCEP 2 mM, Tris 50 mM at pH 8, NaCl 500 mM, glycerol 1%, Protease Inhibitor Cocktail. The lysis extract was centrifuged for 20 min at 25,000 x g, 4°C. Supernatants were loaded onto a HisTrap HP 1 ml column (GE Healthcare) in HEPES 50 mM, KCl 500 mM, NaCl 500 mM, MgCl2 2 mM, TCEP 1 mM, glycerol 2%, mellitic acid 0.002%, Protease Inhibitor Cocktail (Roche), imidazole 5 mM, pH 7.5 and eluted with imidazole. The fractions recovered from the Ni-NTA were further purified by size exclusion chromatography (Superdex 200, for SpoT, SpoT_D81G_, SpoT_cat_, SpoT_ΔACT_ and EI^Ntr^, and Superdex 75 for HPr, EIIA^Ntr^, EIIA^Ntr^_H66E_ and EIIA^Ntr^_H66A_). Purified samples of SpoT were also used to immunize rabbits in order to produce anti-SpoT polyclonal antibodies.

### *In vitro* phosphorylation of EIIA^Ntr^

To phosphorylate EIIA^Ntr^, *B. subtilis* EI, *C. crescentus* HPr and *C. crescentus* EIIA^Ntr^ were mixed at final concentrations of 2.5 *µ*M, 2.5 *µ*M and 50 *µ*M respectively, in phosphorylation buffer [25 mM Tris pH 7.4, 10 mM KCl, 10 mM MgCl2, 1 mM DTT] and incubated at 37 °C for 10 min. The phosphorylation mix was then incubated in 2 mM of PEP at 37 °C for 30 min. Phosphorylated EIIA^Ntr^ (EIIA^Ntr^~P) was further purified by size exclusion chromatography on a Superdex 75 and dialysed for 72 hours at 4°C, in HEPES 50 mM, KCl 500 mM, NaCl 500 mM, MgCl2 2 mM, TCEP 1 mM, glycerol 2% and stored in glycerol 20%, at -20°C.

### *In vitro* hydrolase assay

To assess SpoT hydrolase activity, ^32P^ppGpp was synthesized by enzymatic reaction catalysed by RelA from *Chlorobaculum tepidum*. To synthesize ^32P^ppGpp, 1 *µ*M of RelA_Ctep_ was incubated in TCEP 1 mM, MgCl_2_ 1 mM, NaCl 50 mM, Tris 10 mM at pH 7.4. Then, 100 *µ*M GDP were added to the synthesis reaction and incubated for 10 min at 37 °C, followed by an addition of 3 pM [γ^32^P] ATP (PerkinElmer) incubated for 45 min, 37°C. ^32P^ppGpp was extracted from the reaction medium by centrifugation in Amicon 3K Centrifugal filter (Millipore) at 13,000 x g for 25 min. The hydrolase assays were performed by incubating for 10 min at 37 °C (i) 1 or 2 *µ*M of SpoT1-373 or SpoT_ΔACT_ (**Figure 4a**), (ii) 1 *µ*M of SpoT with increasing concentrations of EIIA^Ntr^, EIIA^Ntr^_H66A_ or EIIA^Ntr^_H66E_ (5, 10 and 20 *µ*M) (**Figure 4b-c**) or (iii) 1 *µ*M of SpoT with 20 *µ*M of purified EIIA^Ntr^~P (**Figure 4 – figure supplement 1**). The reaction mix was then incubated with 10 *µ*l of ^32P^ppGpp to a final volume of 30 *µ*l. The reaction was stopped by adding 2 *μ*l of 12 M formic acid. Reaction products were separated by thin layer chromatography (TLC) by transferring 2 *μ*L of the reaction medium onto a TLC PEI Cellulose F membrane (Millipore). Chromatography membranes were placed in 1 M of KH_2_PO_4_ buffer, pH 3.0, for 50 min at room temperature. The dried TLC membrane was placed in a Phosphor Screen plate (GE Healthcare) for 1 hr and the Phosphor Screen revealed with a phosphoimager.

## Acknowledgements

We are grateful to Emanuele Biondi, Gabriele Klug, Anne Galinier and Josef Deutscher for providing strains and/or plasmids. We thank the members of the BCcD team for critical reading of the manuscript and helpful discussions; Guy Houbeau at the Animal Care Facility of the University of Namur for immunizing rabbits with purified SpoT. This work was supported by a Research Credit (CDR J.0169.16) from the Fonds de la Recherche Scientifique – FNRS to R.H and FNRS-EQP U.N043.17F, FRFS-WELBIO grant (CR-2017S-03), FNRS-PDR (PDR-T.0066.18) to A.G-P, the Programme ‘Actions de Recherche Concertée’ 2016-2021 from the ULB and the Fonds d’Encouragement à la Recherche ULB (FER-ULB) to A.G-P. S.R. was and J.C-M. is holding a FRIA (Fund for Research Training in Industry and Agriculture) fellowship from the Fonds de la Recherche Scientifique – FNRS. R.H. is a Research Associate of the Fonds de la Recherche Scientifique – FNRS.

## Author Contributions

S.R. and R.H conceived and designed the experiments. S.R. performed all the experiments except otherwise stated. A.M. did the cloning and preliminary tests for proteins purification and *in vitro* phosphorylation assays. J.C-M purified the proteins for the biochemical assays and performed the *in vitro* HD assays (Figure 4). S.R., J.C-M, A.G-P and R.H analyzed the data. S.R. and R.H. wrote the paper.

## References

Arenz, S., Abdelshahid, M., Sohmen, D., Payoe, R., Starosta, A.L., Berninghausen, O., Hauryliuk, V., Beckmann, R., and Wilson, D.N. (2016). The stringent factor RelA adopts an open conformation on the ribosome to stimulate ppGpp synthesis. Nucleic Acids Res 44, 6471–6481.

Battesti, A., and Bouveret, E. (2006). Acyl carrier protein/SpoT interaction, the switch linking SpoT-dependent stress response to fatty acid metabolism. Mol Microbiol 62, 1048–1063.

Beaufay, F., Coppine, J., Mayard, A., Laloux, G., De Bolle, X., and Hallez, R. (2015). A NAD-dependent glutamate dehydrogenase coordinates metabolism with cell division in Caulobacter crescentus. EMBO J 34, 1786–1800.

Boutte, C.C., and Crosson, S. (2011). The complex logic of stringent response regulation in Caulobacter crescentus: starvation signalling in an oligotrophic environment. Mol Microbiol 80, 695–714.

Brown, A., Fernandez, I.S., Gordiyenko, Y., and Ramakrishnan, V. (2016). Ribosome-dependent activation of stringent control. Nature 534, 277–280.

Casadaban, M.J., and Cohen, S.N. (1980). Analysis of gene control signals by DNA fusion and cloning in Escherichia coli. J Mol Biol 138, 179–207.

Chiaverotti, T.A., Parker, G., Gallant, J., and Agabian, N. (1981). Conditions that trigger guanosine tetraphosphate accumulation in Caulobacter crescentus. J Bacteriol 145, 1463–1465.

Curtis, P.D., and Brun, Y.V. (2010). Getting in the loop: regulation of development in Caulobacter crescentus. Microbiol Mol Biol Rev 74, 13–41.

De Nisco, N.J., Abo, R.P., Wu, C.M., Penterman, J., and Walker, G.C. (2014). Global analysis of cell cycle gene expression of the legume symbiont Sinorhizobium meliloti. Proc Natl Acad Sci U S A 111, 3217–3224.

Galinier, A., Haiech, J., Kilhoffer, M.C., Jaquinod, M., Stulke, J., Deutscher, J., and Martin-Verstraete, I. (1997). The Bacillus subtilis crh gene encodes a HPr-like protein involved in carbon catabolite repression. Proc Natl Acad Sci U S A 94, 8439–8444.

Gentry, D.R., and Cashel, M. (1996). Mutational analysis of the Escherichia coli spoT gene identifies distinct but overlapping regions involved in ppGpp synthesis and degradation. Mol Microbiol 19, 1373–1384.

Gonzalez, D., and Collier, J. (2014). Effects of (p)ppGpp on the progression of the cell cycle of Caulobacter crescentus. J Bacteriol 196, 2514–2525.

Goodwin, R.A., and Gage, D.J. (2014). Biochemical characterization of a nitrogen-type phosphotransferase system reveals that enzyme EI(Ntr) integrates carbon and nitrogen signaling in Sinorhizobium meliloti. J Bacteriol 196, 1901–1907.

Gropp, M., Strausz, Y., Gross, M., and Glaser, G. (2001). Regulation of Escherichia coli RelA requires oligomerization of the C-terminal domain. J Bacteriol 183, 570–579.

Hallez, R., Delaby, M., Sanselicio, S., and Viollier, P.H. (2017). Hit the right spots: cell cycle control by phosphorylated guanosines in alphaproteobacteria. Nat Rev Microbiol 15, 137–148.

Haseltine, W.A., and Block, R. (1973). Synthesis of guanosine tetra- and pentaphosphate requires the presence of a codon-specific, uncharged transfer ribonucleic acid in the acceptor site of ribosomes. Proc Natl Acad Sci U S A 70, 1564–1568.

Hauryliuk, V., Atkinson, G.C., Murakami, K.S., Tenson, T., and Gerdes, K. (2015). Recent functional insights into the role of (p)ppGpp in bacterial physiology. Nat Rev Microbiol 13, 298–309.

Jain, V., Saleem-Batcha, R., China, A., and Chatterji, D. (2006). Molecular dissection of the mycobacterial stringent response protein Rel. Protein Sci 15, 1449–1464.

Krol, E., and Becker, A. (2011). ppGpp in Sinorhizobium meliloti: biosynthesis in response to sudden nutritional downshifts and modulation of the transcriptome. Mol Microbiol 81, 1233–1254.

Lee, C.R., Park, Y.H., Kim, M., Kim, Y.R., Park, S., Peterkofsky, A., and Seok, Y.J. (2013). Reciprocal regulation of the autophosphorylation of enzyme INtr by glutamine and alpha-ketoglutarate in Escherichia coli. Mol Microbiol 88, 473–485.

Lesley, J.A., and Shapiro, L. (2008). SpoT regulates DnaA stability and initiation of DNA replication in carbon-starved Caulobacter crescentus. J Bacteriol 190, 6867–6880.

Loveland, A.B., Bah, E., Madireddy, R., Zhang, Y., Brilot, A.F., Grigorieff, N., and Korostelev, A.A. (2016). Ribosome*RelA structures reveal the mechanism of stringent response activation. Elife 5.

Mechold, U., Murphy, H., Brown, L., and Cashel, M. (2002). Intramolecular regulation of the opposing (p)ppGpp catalytic activities of Rel(Seq), the Rel/Spo enzyme from Streptococcus equisimilis. J Bacteriol 184, 2878–2888.

Mendrygal, K.E., and Gonzalez, J.E. (2000). Environmental regulation of exopolysaccharide production in Sinorhizobium meliloti. J Bacteriol 182, 599–606.

Potrykus, K., and Cashel, M. (2008). (p)ppGpp: still magical? Annu Rev Microbiol 62, 35–51.

Ronneau, S., Petit, K., De Bolle, X., and Hallez, R. (2016). Phosphotransferase-dependent accumulation of (p)ppGpp in response to glutamine deprivation in Caulobacter crescentus. Nat Commun 7, 11423.

Wells, D.H., and Long, S.R. (2002). The Sinorhizobium meliloti stringent response affects multiple aspects of symbiosis. Mol Microbiol 43, 1115–1127.

